# Valuation of carbon services produced by wild animals finances conservation

**DOI:** 10.1101/2021.10.19.464992

**Authors:** Fabio Berzaghi, Ralph Chami, Thomas Cosimano, Connel Fullenkamp

## Abstract

Filling the global biodiversity financing gap will require significant investments from financial markets, which demand credible valuations of ecosystem services and natural capital. However, current valuation approaches discourage investment in conservation because their results cannot be verified using market-determined prices. Here, we bridge the gap between finance and conservation by valuing only wild animals’ carbon services for which market prices exist. By projecting the future path of carbon service production using a spatially-explicit demographic model, we place a credible value on the carbon-capture services produced by African forest elephants. If elephants were protected, their services would be worth $35.9 billion (24.3-41.2) and store 377 MtC (318-388) across tropical Africa. Our methodology can also place lower bounds on the social cost of nature degradation. Poaching would result in $10-14 billion of lost carbon services. Our methodology enables the integration of animal services into global financial markets with major implications for conservation, local socio-economies, and conservation.

## Introduction

The collapse of biodiversity and ecosystems threatens the long-term sustainability of the biosphere and human society^1^. Large investments from financial markets are needed to protect and restore natural ecosystems. Yet relatively few resources have been committed to protecting nature ^2^. Global financial markets cannot promote significant investment into natural capital until credible valuations of natural resources become widely available. Current valuations use shadow prices, willingness to pay, or other implicit or indirect measurements^3,4^ and are thus disconnected from market prices. This disconnect discourages investors, who rely on market price information, and has created a shortfall in social spending on nature protection known as the “global biodiversity financing gap” ^5^.

Bridging the biodiversity financing gap may require a more modest approach to valuation that focuses only on market-valued services produced by nature. Currently, carbon storage and sequestration produced by species and ecosystems^6–8^ are the only market-valued services that could support investments and trading in the near term. Several national and transnational carbon markets already exist and a global market will likely emerge ^9^. Carbon prices have been steadily rising under increased global demand to reach carbon neutrality. This scenario, however, presumes that carbon services produced by natural entities are sufficiently valuable, and investor interest sufficiently high, to support the development of a market for this instrument.

Mounting evidence shows that wild animals influence carbon fluxes, promote carbon storage^6,7,10^, and should be part of nature-based solutions to mitigate climate change^11^. Other natural entities perform carbon services^4,8^, but valuing and protecting animals inherently involves conserving and restoring their natural habitats. This prevents the “empty forest” effect observed in other CO2 emission reduction schemes centered on habitat carbon services such as UN-REDD+^12^. Habitat-centered schemes do not offer sufficient protection for animals and lead to defaunation that also undermines carbon storage^12^. Investing in animal carbon services would provide a win-win model to preserve ecosystems, reduce biodiversity loss, and mitigate climate change. These added benefits cannot be valued but are nonetheless increasingly important because of the rise of environmental, social, and governance (ESG) investing during the past decade^13^. Financial crisis and increased public pressure for ESG reporting incentivize investments with positive outcomes for the environment and society^14^. The high-ESG rating of our proposed financial framework make it more likely to attract investor.

### Financing conservation through market-valued services

Modern finance provides a framework for establishing markets in natural services through a new class of financial assets that consist of claims on these services. These claims establish the ownership of the services, which may be sold to other individuals, corporations, non-government organizations, and governments. Under this model, governments retain ownership of natural resources and hold the initial claims on the services they produce. Investors could purchase the rights to the flows of services to receive income from them, or in anticipation of capital gains from reselling the rights later. The claims on the services are valueless or decrease in value if the resources producing them are harmed or destroyed. Consequently, investors would require that at least part of the proceeds of the sales must be earmarked for protecting and restoring the resources. In cases where natural entities are granted legal rights^15^, degradation or destruction could also result in fines or penalties, which further motivates service holders to protect natural entities. In this way, the creation of natural asset-backed securities would provide both incentives for preservation of nature as well as the funding mechanism for it. This scheme incentivizes both the public and private sectors to make long-term financial commitments to nature conservation.

### Case study: forest elephant carbon services

Given the demand for carbon services and high-ESG investments, we examine the extent of a market based on animal carbon services through the case study of the African forest elephant (*Loxodonta cyclotis*). Elephants contribute to increasing rainforest aboveground carbon (AGC) by reducing the density of small trees through trampling and consumption^10^. Lower tree density leads to less competition for resources, allows trees to grow larger, and promotes late-succession trees which store more carbon per volume than other type of trees. Overall, forests with elephants store 7-14% more carbon compared to forests without them^10^. Forest elephants are in rapid decline but were once widespread across tropical Africa^16^. Additionally, elephants are a keystone species performing other critical and unique ecological functions such as seed and nutrient dispersal which benefit the whole ecosystem and promote biodiversity^16^. All these conditions make the valuation of elephant services an important proposition for funding their conservation across countries and obtain carbon and additional ecosystem-wide benefits.

We valued the carbon services of elephants in 79 tropical rainforest Protected Areas (PAs) across nine African countries under different population growth scenarios (natural, current poaching, and heavy poaching). Estimates of elephant contribution to aboveground carbon ^10^ were integrated in a spatially-explicit demographic model based on empirical observations ^17^ and considering different forest regeneration rates which could influence carbon gains (Methods). This demographic model permits the quantification of the effect of rebounding elephant populations on the AGC of Protected Areas in the next 100 years. These results were then used in a financial framework to value elephant services and the losses due to poaching. The financial framework evaluated the annual cashflow of carbon capture services produced by current elephants and their contribution to future elephant generations (Methods). We used the carbon price of $51.56 per tCO_2_ based on the EU-ETS market discounted at 2% to calculate the future value of elephant services (Methods). The magnitude of African forest elephants’ contributions to carbon capture are large enough, and the market price of carbon has recently become high enough, to imply that a sizable market for investment in elephant-related carbon credits could be created.

### Elephant populations and Protected Areas

The selected 79 tropical rainforest PAs cover 537,722 km^2^ and host an estimated population of ~99,000 elephants (Fig. 1a, Supplementary Table 1). Most of the PAs (n = 69) are in central African countries: Cameroon (20), Central African Republic (CAR, 3), Democratic Republic of Congo (DRC, 13), Equatorial Guinea (1), Gabon (17), and Republic of the Congo (15). The others are in Nigeria (5), Rwanda (2), and Uganda (3). Current population density varies greatly among PAs (0-0.92 elephants/km^2^, Fig. S1). Protected Areas are mostly National Parks (40) and Natural or Forest Reserves (13). The demographic model predicted that after 100 years without poaching the elephant population would quadruple from ~99,000 to ~394,000 individuals (Fig. S1).

**Fig. 1.**
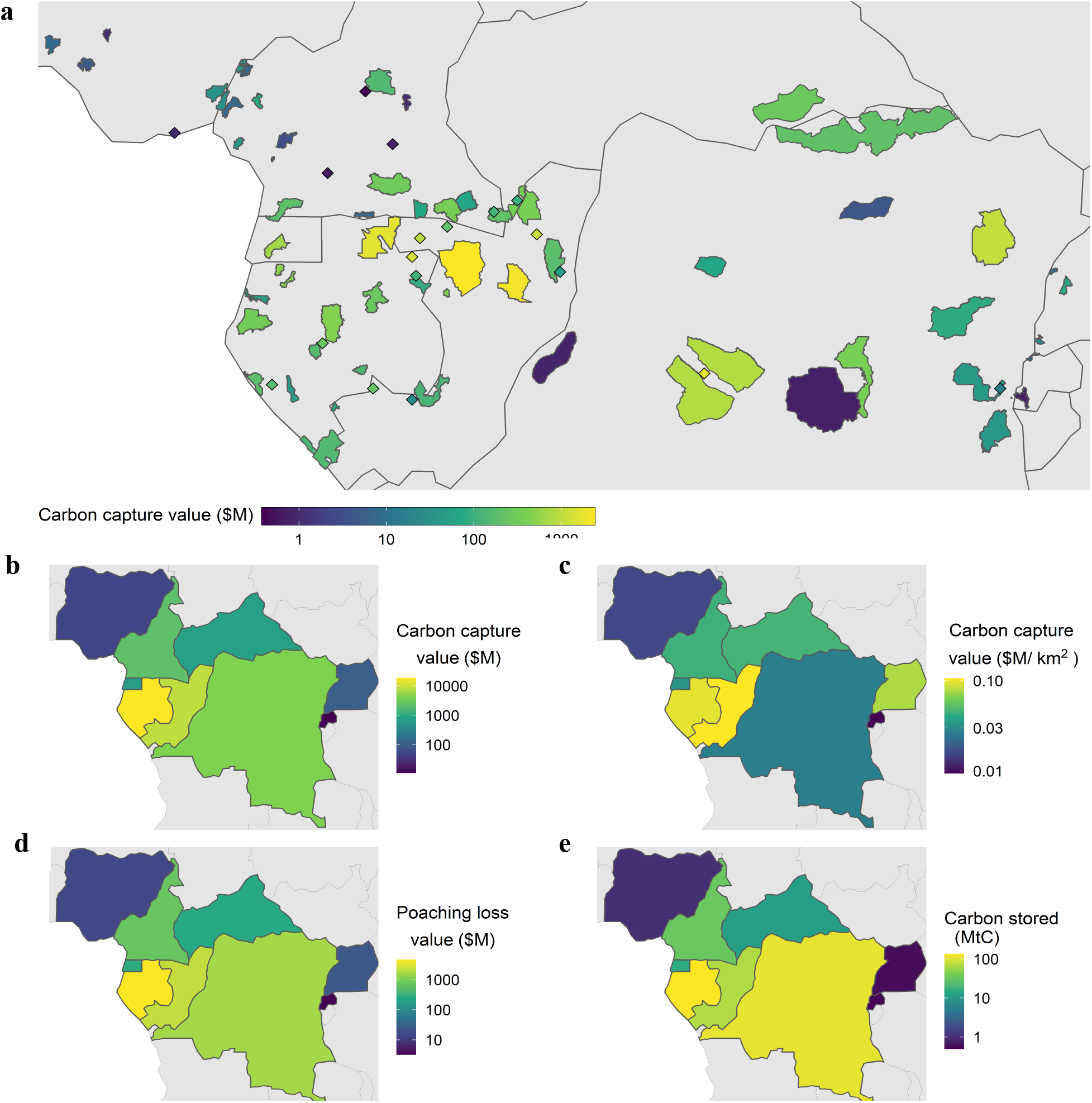
Carbon capture value and stored carbon due to forest elephant activity in 100 years. **a**, Diamond represent PAs not present in the WDPA. The extent of some PAs does not fully match with elephant habitat (see Methods). The PA “Rest of Gabon” is not displayed as it covers ~54% of the country (value $12.8 billion). **b,** Total value of elephant carbon service and **c** per km^2^ value (ratio of total value to total extent of PAs). **d,** Loss of value caused by depressed population growth under current poaching intensity. **e,** Sum of carbon stored across all PAs within each country.

### Carbon sequestration and valuation

The elephant population growth would result in ~377 MtC (318-388) stored in forests due to elephants (~6.2% increase in AGC) and a potential market value of $35.9 billion (24.3-41.2) across all nine countries (Fig. 2 inset). This value does not include the costs of protecting forests and elephants, helping local communities with human-wildlife conflicts, and implementing the conservation program. These results assume that anthropogenic disturbances such as deforestation or degradation would be minimal within PAs. The value of services in individual countries varies widely between $11-18,800 million because of differences in present-day population and extent of PAs (Fig. 1b, Fig. 2, Supplementary Table 1). Nigeria, Rwanda, and Uganda are at the edges of the forest elephant range and have a few small PAs (Fig. 1a, Fig. 3). Uganda has however higher chances of approaching its maximum carbon-sink potential given its high elephant density (Fig. 1b, 3a). Gabon and the Republic of Congo are also well positioned to optimize elephant services due to their large PAs and more abundant populations (Fig. 1b, 3a). The other countries have extended PAs but small populations; in particular the DRC (second largest extent of PAs, Fig.3). This reduces their potential for carbon capture and value (Fig. 3), which is nonetheless still the third largest. The negative effect of small populations is also observed in the carbon capture value per km^2^ which is low in DRC and CAR compared to the potential offered but their large PAs (Fig. 1c). Small current populations result in unrealized carbon value as it takes longer to restore populations and fully benefit from their services (Fig. 3a). Small PAs limit the country-level value of carbon services because elephant range is restricted. Extending PAs could be a solution if matched with increased protection to avoid poaching and reduction in elephant-human conflict.

**Fig. 2.**
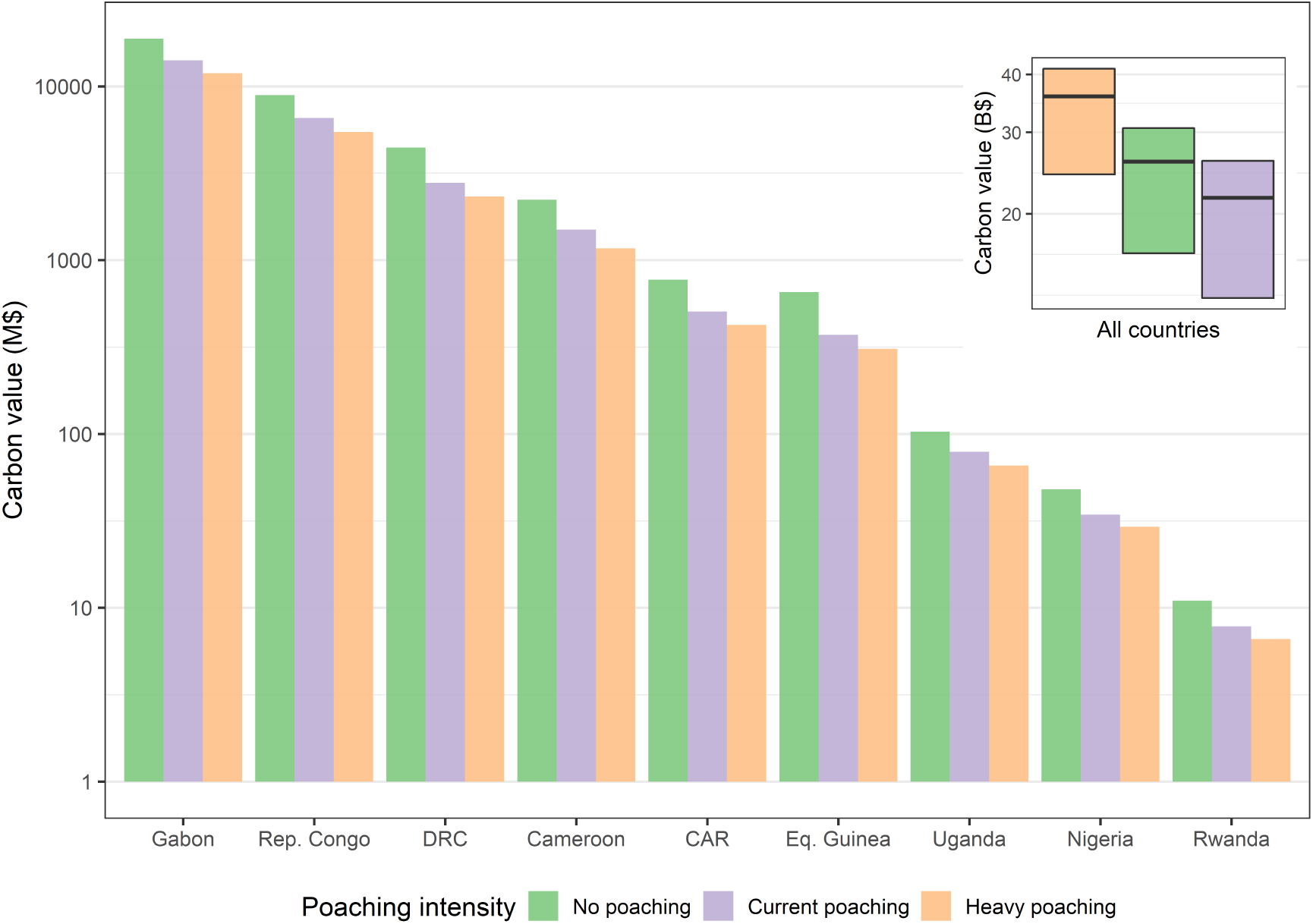
Value of forest elephant services in African countries under different poaching intensities. Values are the cumulative sum of yearly carbon service in PAs over an investment horizon of 100 years and include the contribution of present and future generations of elephants. The inset shows the total of all countries with upper and lower bound calculated with a, respectively, faster and slower forest regeneration rate compared to the base scenario (see Methods).

**Fig. 3.**
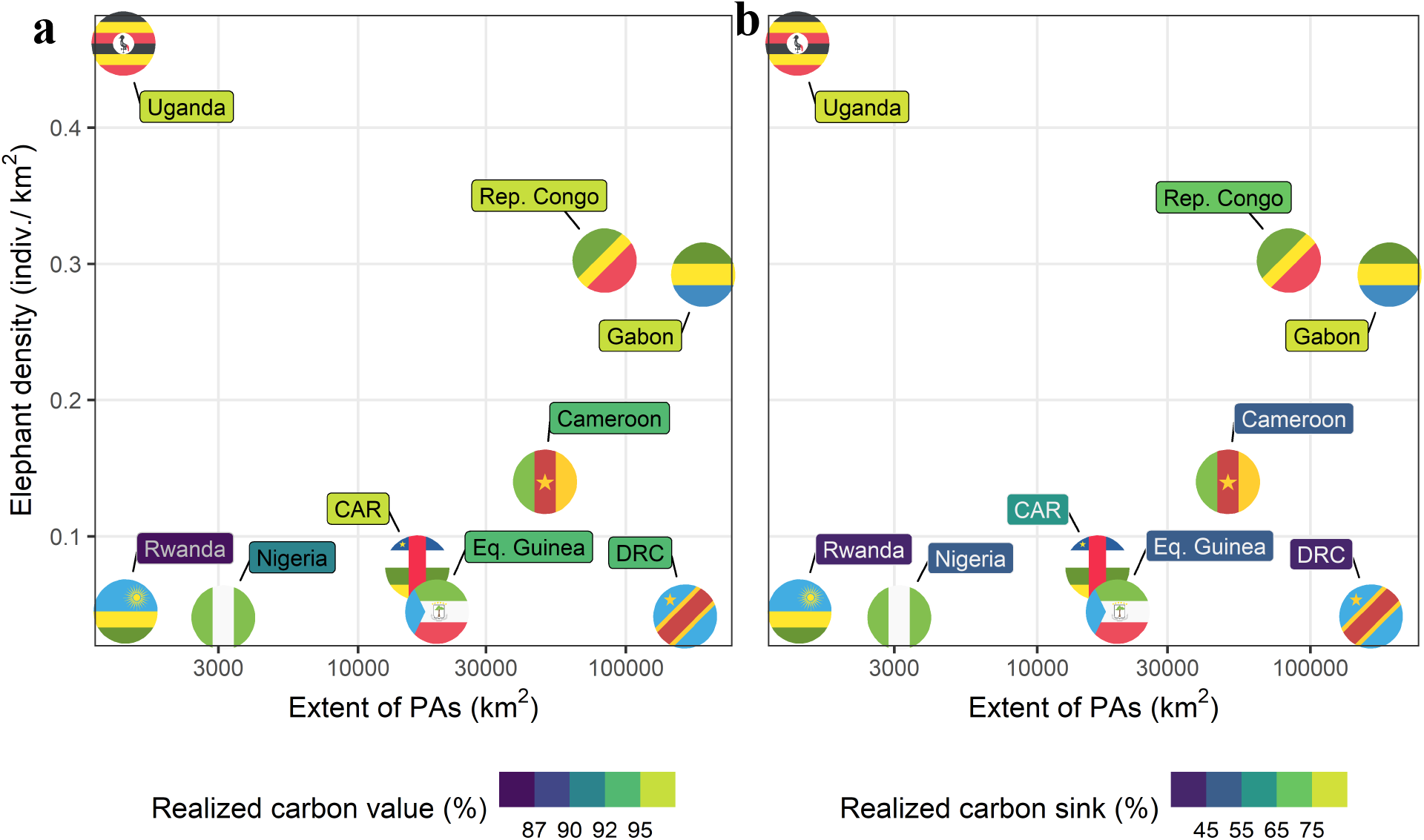
Carbon value and carbon sink potential attained by countries within the next 100 years. The percentage represents the fraction of **a** value or **b** AGC stored under no poaching compared to the maximum value if elephant populations returned to their natural density of 1 elephant/km^2^.

### The cost of poaching

In 100 years, current and heavy poaching rates ^17^ (Methods) would limit population growth to ~283,000 and ~191,000, respectively (Fig. S1). Consequently, the value of carbon services is severely reduced by $10 billion (27.7%) and $14 billion (39.7%), respectively, compared to the no-poaching scenario (Fig. 2). High losses due to poaching are observed in Gabon, DRC, and Congo and intermediate losses in CAR, Cameroon, and Equatorial Guinea (Fig. 1d). The losses incurred in the poaching scenarios are initially small but after 30-50 years they already amount to $6-11 billion (Fig. S2). Once most the carbon-capturing potential is achieved, long-term losses would be ~$10 billion or more depending on uncertainty in the time needed for AGC to reach equilibrium with elephants (Fig. S2 and Methods) and differences in local poaching rates. After 100 years, the bulk of the carbon sink is realized in Gabon, Congo, and Uganda (Fig. 1e and 3b). Instead, the other countries would attain only between 37-55% of their potential carbon sequestration (Fig. 3b). All nine countries have small carbon footprints implying that elephants could further expand their role as carbon sinks for several years. A maximum potential carbon sink of 634 MtC would only be accrued over a much longer time window (200-350 years) as current populations in most PAs are small (Supplementary Table 1).

### Creating markets around conservation

To contextualize these calculations in terms of single PAs, consider Nouabale-Ndoki National Park (Republic of Congo) hosting ~1800 elephants at a density of 0.45. The carbon capture services provided by these elephants would be worth ~$380 million. This amount is comparable to the average market capitalization of a publicly traded company in the Russell 2000 index, a measure of 2,000 U.S. small stocks having a median value of slightly less than $1 billion. This comparison implies that the size of the Nouabale-Ndoki market alone would be large enough to attract institutional investors. These investors include pension funds, endowments, mutual funds, and hedge funds. In addition, millions of individual investors, for whom the trading of equity is an everyday occurrence, would also participate in the Nouabale-Ndoki market.

The above calculations show that the carbon capture services of African forest elephants could form the basis of an investment market worth over $35 billion. Our carbon price is market-based and in line with IPCC reports and literature^18,19^ but increases in the price and its variation across regions^18^ might also affect these kind of valuations. Forest elephant services are only one part of a potentially much larger global market in animal carbon services. Creating these markets would involve the associated costs for the protection (e.g., anti-poaching activities, creation of PAs), species reintroduction and restoration of ecosystems (e.g., rewilding), and compensations for lost income for local communities and their direct involvement throughout the project (e.g., policy, implementation, and management)^20^. Added benefits of restoring and preserving ecosystems include their increased resilience to future climate disturbances^21^ which threaten ecosystems’ health and services. Researchers are establishing that many other animals including marine and terrestrial vertebrates^6,7^ and invertebrates^6^ play important roles in carbon cycling. The total market value of carbon services may be measured in the trillions of dollars—smaller than global equity markets, but as large as the markets for important types of bonds such as commercial paper. More broadly, the techniques used in this paper can be applied to any animal service that can be measured, and to which market prices may be assigned. Animal services, therefore, represent an entire asset class whose market potential may rival that of existing financial instruments.

### Challenges and opportunities

The obvious next question is how to develop these markets as quickly as possible, in order to enable actual investments to fund preservation and restoration of vanishing species and habitats. Many steps are involved in financial market development^22^. Certification of the carbon sequestration produced by forest elephants (and other species) is necessary for investor acceptance of financial instruments based on this service. The best approach is to start small, with a demonstration case in a few PAs with a good record of elephant protection and intact habitat. Protected areas should be kept intact as much as possible before and while elephant populations are recovering and synergies with other climate change mitigation strategies could further preserve biodiversity and carbon stocks^23^. Collaboration between governments, including local communities, and institutional investor representatives over the design of the financial instrument will greatly increase the likelihood of successful issuance. If certification and instrument design obstacles can be overcome, the outlook is positive. The increasing global demand for carbon offsets, driven by corporate and government pledges to reach carbon neutrality, and high-ESG portfolios present an unprecedented opportunity to develop financial markets that support conservation, backed by the services produced by the nature being protected.

## Methods

### Carbon Capture Services of African Forest Elephants

African forest elephants (*Loxodonta cyclotis*) facilitate the capture of large quantities of carbon through different mechanisms. First, the average body mass of a mature forest elephant contains 720 kg of carbon ^24^. After death, the carbon contained in bodies is mostly released back into the atmosphere in the form of CO_2_. However, a stable population of elephants will continually store carbon in proportion to the number of individuals. Any increase to the stable population implies that additional carbon is captured and stored in elephant bodies.

Most importantly, African forest elephants facilitate carbon sequestration through their effect on the forest ecosystem. While moving through the forest and foraging for food, elephants reduce the density of trees smaller than 30 cm in diameter. This reduction in tree density changes light and water availability in the forest, leading to an increase in the proportion and the average size of late-succession trees. Late-successional are slow-growing, canopy-dominant trees with a higher carbon density (kgC/m^3^) compared to other tree types. As late-successional trees become larger and more abundant, there is a net increase in the forest aboveground carbon due to elephant activity ^10^.

Projecting the future population of elephants is essential to estimating the full value of the carbon capture services they produce. The quantities of services produced are proportional to population, and current populations are much smaller than their natural pre-industrial levels, due to poaching and habitat loss. The population is currently estimated at less than 100,000, compared to more than 1,000,000 individuals before widespread poaching ^25^. Thus, we initially define a demographic model to project future changes in elephant populations under different poaching scenarios. The demographic model is than used to calculate the value of carbon in elephant bodies and of their carbon-capturing services.

### Elephant population model

We forecast population growth using a logistic model under three scenarios: 1) natural mortality under no poaching, 2) mortality under current poaching rates, and 3) mortality under high poaching rates (see below for further details on mortality rates). The logistic model estimated the evolution of the forest elephant populations in Protected Areas (PAs) across Africa. The annual growth rate of elephants is given by

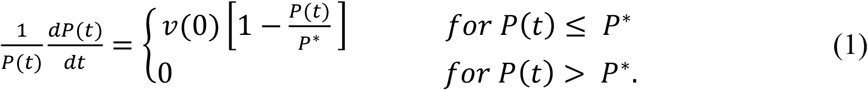

Where *P(t)* is the population at time *t* and *P*\* is the population carrying capacity estimated at 1 elephant/km^2^ based on literature. The left-hand side of eq. 1 is the percentage change in the population of elephants. The right-hand side of eq. 1 indicates that when *P(t)* is less or equal to *P*\* the growth rate starts at 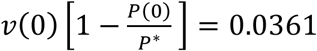 in the natural mortality scenario and 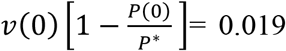 in the current-poaching scenario. These population growth rates were determined following the only long-term study of forest elephant demography ^17^. We added a high-poaching scenario where the initial growth rate is 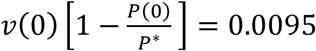 which is half the growth rate under poaching reported in ^17^. This additional scenario was needed to account for the high variability in poaching rates across central Africa. The 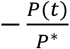 term represents a decrease in the growth rate of elephants as the population approaches carrying capacity. The initial growth rate at *P(0)* decreases towards zero until *P(t)* = *P*\*.

The solution to equation (1) as *t* goes from 0 to infinity follows ^26,27^:

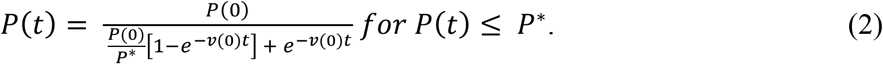

The initial growth rate of the population under no poaching is determined by

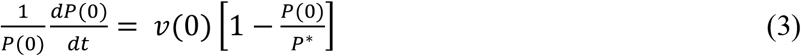

The initial population *P*(0) and the population at carrying-capacity *P*\* are required to solve equation (1) for *P* at each iteration. These parameters were determined for each PAs following the African Elephant Database ^28^ (AED) and literature ^29^. We use the term PA in a broader sense as the AED identifies “Input zones” which are sometime outside PAs. Further, because the AED does not distinguish between forest and savanna populations, we retained only the protected areas covered by tropical rainforests within potential forest elephant range. We excluded forest concessions, mountainous areas, and the savanna part of mixed-vegetation PAs. These data were used to determine current population density, potential range (i.e., PA extent), and average above ground carbon (Supplementary Table 1). Population density at equilibrium was set at 1 elephant/km^2^.

### Value of carbon capture in elephant bodies

The value of carbon stored in elephant bodies is marginal compared to the carbon captured by elephants through their interactions with the forest. We performed the calculations for completeness and because it might be of interest for species attaining larger total population biomass such as ocean vertebrates ^7^. We estimate that an average elephant weighs 3000 kg of which 24% is carbon ^24^. The carbon sequestered in the body is multiplied by 3.667 to obtain its CO_2_ equivalent (*C_b_*).

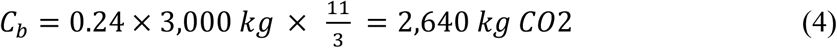

The value of carbon sequestered per elephant body is calculated by multiplying *C_b_* by the price of CO_2_ per kg × 10^3^ (*C_p_*). The average CO_2_ price reported in the European Union Emissions Trading System market in 2021 was $51.56 ^30^. This price is the average over the period January 1, 2021 to June 10, 2021 using historical futures prices: European Climate Exchange EU allowance Futures, Continuous Contract #1. One EU allowance gives the holder the right to emit one ton of CO_2_. This price is converted from Euros to US dollars using the average US/Euro exchange rate over the same time period from the St. Louis Federal Reserve Bank. The yearly value of CO_2_ captured in the population is equal to the increase in population multiplied by the CO_2_ captured per elephant multiplied by *C_p_* so that the market value for this service during period *t* + *i* is given by

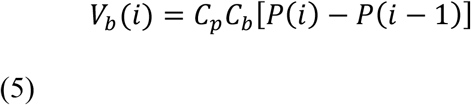

To find the present value of current and future carbon capture, we must assume a discount rate (*d*), which is the return on $1 after one year. Present value of a future cashflow of $1 is discounted by an interest rate, *d*, by 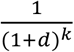. Here, *k* is the numbers of years into the future. This procedure identifies the amount of money needed today equivalent to $1 in *k* years into the future. Once, the $ value is placed into the same time period, all the future values can be added together. We chose a 2% discount rate reflecting both market evidence and the practices in the existing literature ^31,32^. Using *d* = 0.02, the present value of carbon content in the body of all elephants in a PA is:

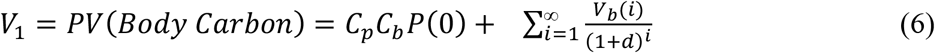

### Value of carbon capture enhancement through interaction with tropical forest

The enhancement of aboveground carbon (AGC) triggered by elephants is determined by their population density and the state of AGC in relation to the equilibrium-AGC with elephants ^10^. Here we assume that PAs are not subject to intense anthropogenic disturbances which might perturb carbon cycling. For example, we excluded areas that are selectively logged. In our case, at a density of 1 elephant/km^2^, equilibrium-AGC is 14% higher compared to AGC in a forest without elephants. As elephants are removed from the system, AGC starts to decrease until it reaches a new equilibrium relative to a lower elephant density. The time to transition from one equilibrium-AGC to another depends on the rate of population decline and the mortality rate of trees. These two rates are needed to estimate how much of the 14% gain has been lost since their decline and to calculate the future contribution of elephants to AGC. Once the potential gain (14% – % lost since decline) is determined, the equilibrium-AGC at a density of 1 elephant/km^2^ can be estimated from the current AGC in each PA. Historical rates of population decline were not available for most of the PAs in our study. Instead, we used the current population density in each PA as an indication of years since decline. The majority of populations across central Africa declined between 20 and 100 years ago with some exceptions of areas afflicted more recently by intensive poaching. A density close to zero might suggest that local populations declined 100 years ago or more. At higher population densities, the years-since-decline would be less. Following these assumptions, we use a linear function to determine the years since decline (*t_d_*) as a function of current population density for each PA. We acknowledge that local declines might not follow linear patterns.

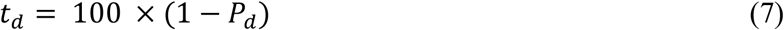

Where *P_d_* (elephants/km^2^) is the population density in a particular PA.

Tree morality rate provides an indication of how fast a forest regenerates itself. Observed mortality rates in tropical forests are highly variable (2-6% per year) and are affected by drought and extreme climatic events ^33,34^. Observed mortality rates of African tropical forests suggest rates between 1% and 2% ^35,36^. However, estimating mortality rates requires long-term studies which are limited in Africa compared to other tropical areas ^35^. We account for this uncertainty by setting mortality rate (*m*_r_) at 1.5% and by performing a sensitivity analysis on this parameter which we use to generate confidence intervals for our results. A yearly mortality of 1.5% implies that 67 years are needed to replace most adult trees with new recruits. Under these conditions, where elephant density is close to zero, *t_d_* would be approaching 100 years. Most adult trees would have been replaced and AGC likely reached its equilibrium without elephants so the potential future gain in AGC would be around 14%. We used *t_d_* and *m*_r_ to calculate the equilibrium-AGC if elephants would return to their original density of 1 elephant/km^2^. The three study cases used for the sensitivity analyses are further explained in the Supplementary Text.

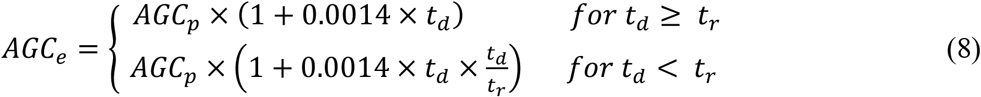

Where *AGC_e_* (10^3^ kgC) is the equilibrium-AGC with elephants, *AGC_p_* (10^3^ kgC) is the present-day AGC, and *t_r_* (years) represents the forest regeneration time calculated as 100/*m*_r_. *AGC_p_* was calculated as the average AGC within each PA according to the boundaries indicated by the United Nations World Database on Protected Areas ^37^ and the most recent AGC map ^38^.

When elephant density starts to increase, the time to reach *AGC_e_* will depend on various factors: the number of years needed to reach 1 elephant/km^2^ (*t_e_*), the spatial heterogeneity of elephant density, and *m_t_*. Because in the majority of PAs *t_e_* is 1.5-5 times larger than *m_t_*, we conservatively use *t_e_* as an indication of the time taken to reach *AGC_e_*. In all cases, the elephants’ contribution to AGC is maximized only when the population reaches carrying capacity, consequently at least *t_e_* years are needed for this process to complete. We assume that AGC increases at a constant rate irrespective of the initial *AGC_p_*. Therefore, we calculate the yearly rate of change in AGC attributable to elephants with the following:

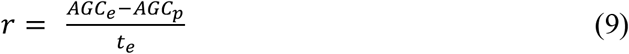

We assume that *r* is constant within each PA. This implies that, even though population density might vary spatially within a PA, an average increase in AGC/km^2^ per year is applied in areas that are reclaimed by elephants. As populations grow, their density will homogenize across the landscape and their effect on AGC will converge to *r*. The average value of *r* across PAs is 0.0885 × 10^3^ kgC/year (s.d. 0.03), which is a small fraction compared to net growth rates of AGC observed in African rainforest 0.6-4 × 10^3^ kgC/year. Our estimated *r* is thus conservative compared to previously published estimates of elephant contribution to AGC. We use *r* and yearly changes in population to calculate the value of carbon capture provided by elephants. The calculation is as follows. At time 1 there is an increase in population of P(1) – P(0), following the population growth model (eq. 2). This new generation enters a plot of forest with *AGC_p_* and increases it to *AGC_e_* over *t_e_*. The size of the plot is adjusted so that the density of elephants in the forest is maintained constant throughout the PA. At time 2, a new generation of elephants is born with size P(2) – P(1), which occupies a new plot and contributes to the growth of AGC as described above. We repeat this process for 1000 generations to ensure convergence of the elephant population to its steady state, at which point the total increase in carbon capture converges to zero. The present value of the carbon capture by current and future generations of elephants has three components given by:

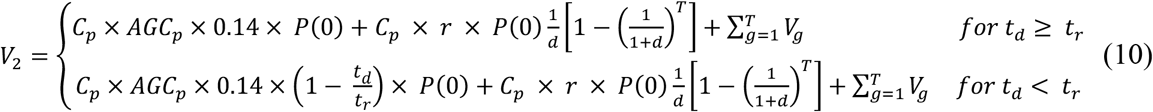

Here, *T* is the investment horizon of the investor, which is the number of years the investor expects to receive payments from the investment. Let *g* be the number of generations, and *V_g_* be the value of each generation contribution to carbon capture given by

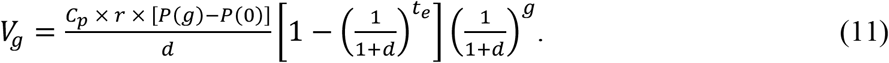

All other variables were defined previously. The first term in eq. 10 is the 14% contribution to AGC by the current population of elephants, P(0), multiplied by the price of carbon credits, *C_p_*. If *t_d_ ≥ t_r_*, the current population is equal to the population density times the area of the PA. Otherwise, we assume that the current population has realized its contribution to *AGC_p_*. In the supplementary appendix the choice between the first and second line in eq. 10 is determined by the mortality of the forest and the initial density of the PA. For example, when the mortality is 1.5% per year, the first line is relevant for the initial density less than or equal to 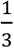. In addition, 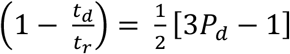 for an initial density greater than 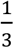, so that the current population does not contribute to the present value of the elephants in the PA, when the initial density is 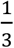. However, the contribution from future generations is highest, since 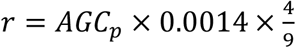 for 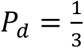 and converges to zero as the initial density of the PA approaches 0.

The next two terms in eq. 10 include the contribution of future generations, *P*(*g*). The population in each generation can be decomposed into a contribution from the initial population, P(0), to the current generation, and the change in the population from the initial population, [*P*(*g*) *− P*(0)], i.e., *P*(*g*) = *P*(0) + *P*(*g*) *− P*(0). The first part P(0) adds for each time the change in the AGC times the market price of carbon capture, so that the second term in eq. 10 is *C_p_* × *r* × *P*(0). This leads to the second contribution in eq. 10. This contribution is multiplied by the present value of an annuity at discount rate, *d*, which pays this contribution for the investment horizon, 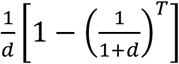. An annuity is a financial contract that pays the same amount each year for a fixed number of years given a discount rate *d*. In the calculations, we use an investment horizon of 100 years. Adding additional years does not have a significant impact on the valuation in eq. 10 because the discount rate lowers the valuation over longer investment horizons. The second part of the contribution from the future generation is the change in the population from the initial population, [*P*(*g*) − *P*(0)], which impacts the population for *t_e_* years by the change in AGC, r multiplied by the price of carbon credits. This third contribution in eq. 11 is multiplied by the present value of an annuity, which pays this contribution for *t_e_* years, 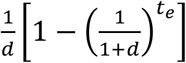. Generation g contributes from time g for *t_e_* years into the future, so we must discount this benefit by 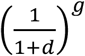 to determine the present value of generation g in eq. 11. Consequently, the present value of each generation *g*’s contribution to carbon capture in eq. 11 is added in the last term of eq. 10 for each generation 1 to T, which is denoted 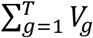. This leads to equation eq. 10 which adds together the present value of the three components of the contributions of current and future generations of elephants. The total value of the elephants is equal to *V* = *V_1_* + *V_2_* following eq. 6 and eq. 10, respectively.

## Data availability

Data for each Protected Area included in the study can be found in the Supplementary Table 1. The code for the ensemble mode is available at https://github.com/fabeit/elephant-valuation.

## Acknowledgments

We thank Stephen Blake, Philippe Ciais, and Dinah Nieburg for providing feedback on the manuscript, and Marie Fauvet for assisting with the calculation of carbon in elephant body.

## Author contributions

Conceptualization: FB, TC, CF, RC

Methodology: lead by FB and TC with contributions from CF and RC

Visualization: FB

Funding acquisition: FB

Writing: FB, TC, CF, RC

## Supplementary Information

is available for this paper

## Materials & Correspondence

Fabio Berzaghi

