## Supplementary information for "Valuation of carbon services produced by wild animals finances conservation"

**This PDF file includes:**

Figs. S1 to S2

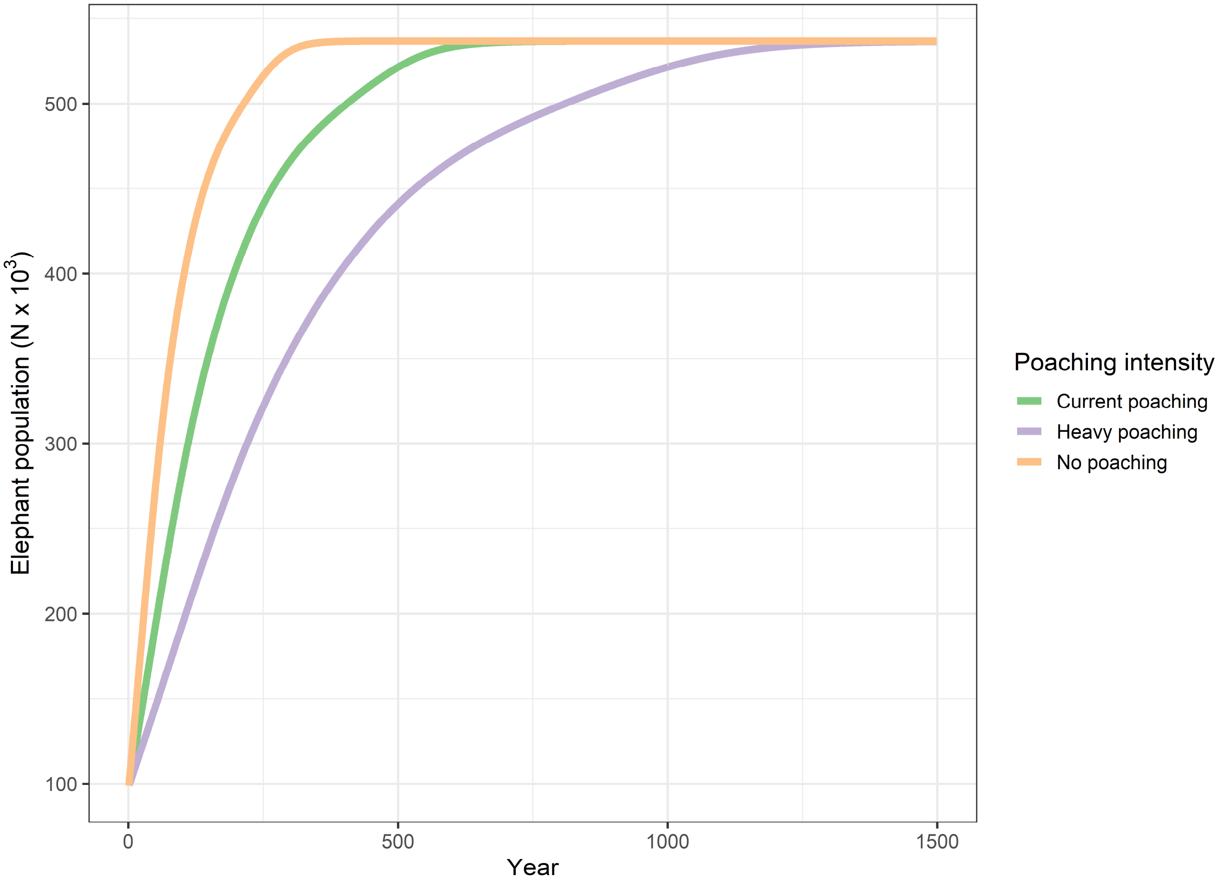


**Fig. S1. Elephant population growth under different poaching intensities.** The year indicates the time elapsed since the start of the simulation.

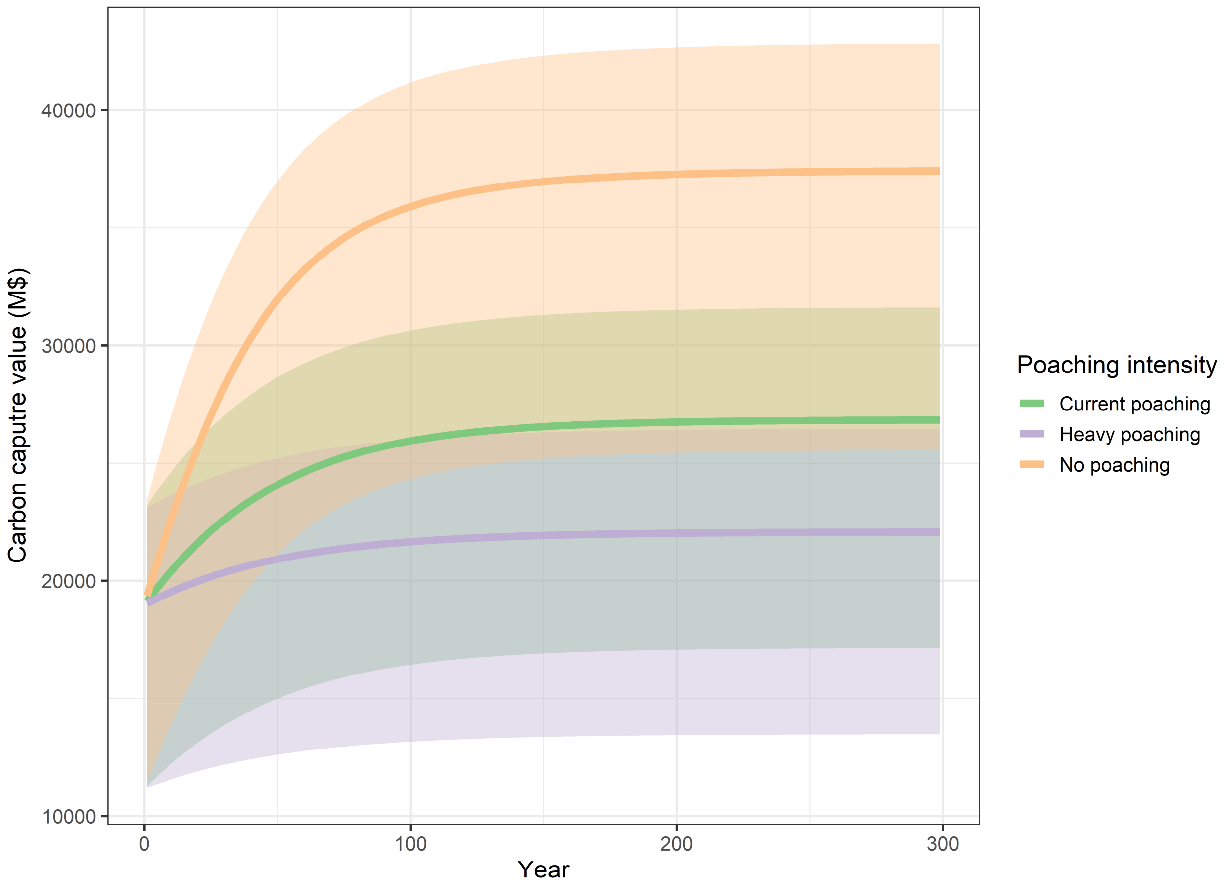


**Fig. S2. Value of the carbon-capturing service of forest elephants under different poaching intensities.** The cumulative value is the sum of all nine countries included in the study and the year indicates the time horizon investment.
