## Supplementary text for "Valuation of carbon services produced by wild animals finances conservation"

**This PDF file includes:**

Supplementary Text

Supplementary Table 1 caption

**Supplementary Text**

Further explanation of carbon capture valuation study cases

Based on the set up in eq. 6-9, eq. 10 (Methods) has three cases dependent on the regeneration rate of African tropical forests. This supplement text explains the specification of eq. 10 for each case.

Case 1. Let $m_r$= 1%, so that 100 years are needed to replace most trees.

Eq. 7 is given by $t_d = 100 \times (1 - P_d)$ so that $t_d \in [100,0]$ for $P_d \in [0,1]$. In this case $t_r = 100$. Consider the first line in eq. 10. Which occurs when $t_d \geq t_r$, so that $100 \times (1 - P_d) \geq 100 \Longrightarrow P_d \leq 0$. This result is false for all PAs, so only the second line in eq. 10 is used in this case. The condition $t_d < t_r$ occurs for $P_d > 0$. As a result, $\left(1 - \frac{t_d}{t_r}\right) = \left(1 - \frac{100 \times (1-P_d)}{100}\right) = P_d$.

By eq. 8 $AGC_e$=$AGC_p \times (1 + 0.14 \times (1 - P_d)^2)$, since $P_d > 0$. In this case $AGC_e = AGC_p \times (1 + 0.14)$ for $P_d = 0$, and $AGC_e = AGC_p$for $P_d = 1$.

By eq. 9 $r = \frac{AGC_p \times (1+0.14 \times (1-P_d)^2) - AGC_p}{100} = AGC_p \times 0.0014 \times (1 - P_d)^2$, so that $r = 0$ for $P_d = 1$ and $r = AGC_p \times 0.0014$ for for $P_d = 0$.

Thus, the present value of elephants given by the second line of eq. 10 becomes

$$C_p \times AGC_p \times 0.14 \times P_d \times P(0) + C_p \times r \times P(0)\frac{1}{d}\left[1 - \left(\frac{1}{1+d}\right)^T\right] + \sum_{g=1}^{T} V_g.$$

For $P_d = 0$ the first term is zero, so there is no contribution from the current population to the present value of the elephants. However, the second and third terms are maximized, since $r = AGC_p \times 0.0014$. This means that all the contribution to the value of the elephants comes from carbon credits.

Case 2. $m_r$= 1.5%, so that 67 years are needed to replace most trees.

In this case $t_r = 67$. Consider the first line in eq. 10. Which occurs when $t_d \geq t_r$, so that $100 \times (1 - P_d) \geq 67 \Rightarrow P_d \leq \frac{1}{3}$. As a result, the first line is true for the lower density PAs. However, the second line in eq. 10 is relevant for $P_d > \frac{1}{3}$. The condition $t_d < t_r$ occurs for $P_d > \frac{1}{3}$. As a result, $\left(1 - \frac{t_d}{t_r}\right) = \left(1 - \frac{100 \times (1-P_d)}{67}\right) = \frac{3}{2} P_d - \frac{1}{2}$.

By eq. 8 $AGC_e$=$AGC_p \times (1 + 0.14 \times (1 - P_d)^2)$ for $P_d > \frac{1}{3}$. In this case, $AGC_e = AGC_p \times (1 + 0.14 \times \frac{4}{9})$ for $P_d = 1/3$, and $AGC_e = AGC_p$for $P_d = 1$.

By eq. 9 $r = AGC_p \times 0.0014 \times (1 - P_d)^2$ for $P_d > \frac{1}{3}$, so that $r = AGC_p \times 0.0014 \times \frac{4}{9}$ for $P_d = \frac{1}{3}$, and $r = 0$ for for $P_d = 1$.

Thus, the present value of elephants given by eq. 10 becomes

$$V_2 = \begin{cases} C_p \times AGC_p \times 0.14 \times P(0) + C_p \times r \times P(0)\frac{1}{d}\left[1 - \left(\frac{1}{1+d}\right)^T\right] + \sum_{g=1}^{T} V_g & for\ P_d \leq \frac{1}{3} \\ C_p \times AGC_p \times 0.14 \times \frac{1}{2}[3\,P_d - 1] \times P(0) + C_p \times r \times P(0)\frac{1}{d}\left[1 - \left(\frac{1}{1+d}\right)^T\right] + \sum_{g=1}^{T} V_g & for\ P_d > \frac{1}{3} \end{cases}$$

For $P_d = 1/3$ the first term in the second line is zero, so there is no contribution from the current population to the carbon sequestration of the forest. However, the second and third terms are maximized, since $r = AGC_p \times 0.0014 \times \frac{4}{9}$. This means that all the contribution to the value of the elephants comes from carbon credits.

<u>Case 3. $m_r$ = 2%, so that 50 years are needed to replace most trees.</u>

In this case $t_r = 50$. Consider the first line in eq. 10. Which occurs when $t_d \geq t_r$, so that $100 \times (1 - P_d) \geq 50 \Rightarrow P_d \leq \frac{1}{2}$. As a result, the first line is true for the lower density PAs.

However, the second line in eq. 10 is relevant for $P_d > \frac{1}{2}$. The condition $t_d < t_r$ occurs for $P_d > \frac{1}{2}$. As a result, $\left(1 - \frac{t_d}{t_r}\right) = \left(1 - \frac{100 \times (1-P_d)}{50}\right) = 2\,P_d - 1$.

By eq. 8 $AGC_e$=$AGC_p \times (1 + 0.14 \times (1 - P_d)^2)$ for $P_d > \frac{1}{2}$. In this case, $AGC_e = AGC_p \times (1 + 0.14 \times \frac{1}{4})$ for $P_d = \frac{1}{2}$, and $AGC_e = AGC_p$for $P_d = 1$.

By eq. 9 $r = AGC_p \times 0.0014 \times (1 - P_d)^2$ for $P_d > \frac{1}{2}$, so that $r = AGC_p \times 0.0014 \times \frac{1}{4}$ for $P_d = \frac{1}{2}$, and $r = 0$ for $P_d = 1$.

Thus, the present value of elephants given by eq. 10 becomes

$$V_2 = \begin{cases} C_p \times AGC_p \times 0.14 \times P(0) + C_p \times r \times P(0)\frac{1}{d}\left[1 - \left(\frac{1}{1+d}\right)^T\right] + \sum_{g=1}^{T} V_g & for\ P_d \leq \frac{1}{2} \\ C_p \times AGC_p \times 0.14 \times [2P_d - 1] \times P(0) + C_p \times r \times P(0)\frac{1}{d}\left[1 - \left(\frac{1}{1+d}\right)^T\right] + \sum_{g=1}^{T} V_g & for\ P_d > \frac{1}{2} \end{cases}$$

**Supplementary Table 1** in file Supplementary_Table1_HFT_classification.xlsx

Properties of Protected Areas included in this study and their valuation under different elephant growth scenarios. The “value” columns express the $ value of forest elephant services after 100 years or their maximum theoretical value. The “carbon” columns express the carbon stored in forests in MtC as a result of elephant population growth. All other columns are self-explanatory and their units indicated within parenthesis.
